# Subcortical and cortical tracking of communication sound envelopes in challenging listening conditions

**DOI:** 10.1101/2022.02.10.479939

**Authors:** S. Souffi, L. Varnet, M. Zaidi, B. Bathellier, C. Huetz, J.-M. Edeline

**Author notes:** Corresponding Author*: Jean-Marc Edeline, UMR 9197, Neuro-PSI (Institut des Neurosciences Paris-Saclay), CNRS - Université Paris-Saclay, Centre CEA Saclay, Bâtiment 151, 91400 Saclay cedex, France. Edmond and Lily Safra Center for Brain Sciences, The Hebrew University of Jerusalem, Edmond J. Safra Campus, Givat Ram, Jerusalem 9190401, Israel.

## Abstract

Humans and animals constantly face challenging acoustic environments such as various background noises restricting the detection, discrimination and identification of behaviorally salient sounds. Here, we disentangled the role of temporal envelope tracking on the decrease in neuronal and behavioral discrimination between communication sounds in situations of acoustic degradations. We simulated responses of auditory nerve fibers and recorded neuronal activity in cochlear nucleus, inferior colliculus, thalamus and auditory cortex in anesthetized guinea-pigs. Furthermore, a Go/No-Go sound discrimination task involving two of the guinea-pig whistles was performed on mice in silence and noise. For all conditions, we found that auditory neurons better track the slow amplitude modulations (<20 Hz) of the stimulus envelopes than the faster ones. In addition, the decrease in neuronal and behavioral discrimination performance in noise can be explained by an increased similarity of the vocalization envelopes in the low frequency range (<20 Hz). Together, these results suggest that slow envelope tracking is a general property of auditory neurons, and any difference between the slow envelopes of natural stimuli allows coping with degraded conditions.

## Introduction

In humans, speech signals are characterized by rhythmic streams of amplitude and frequency modulations (AM and FM) that convey phoneme, syllable, word, and phrase information (Rosen 1992, Varnet et al., 2017, Ding et al., 2017). It is known for several decades that the modulations of slow temporal envelope carry essential cues for speech perception (Drullman et al., 1994; Shannon et al., 1995; Zeng et al., 2005): even in challenging conditions, the human auditory system has the capacity to process highly degraded speech as long as the temporal envelope modulations below 20 Hz are preserved (Drullman et al., 1994a, b; Shannon et al., 1995). Furthermore, the extent to which speech envelopes are preserved by a given transmission system has proved to be a reliable predictor of intelligibility in a variety of listening situations, including noise, fast-acting amplitude compression, or reverberation (Houtgast and Steeneken, 1985; Plomp, 1988; Steeneken and Houtgast, 1980). This is consistent with electroencephalogram (EEG) and magnetoencephalography (MEG) studies in which cortical responses were found in phase with the temporal envelope of speech signals (Ahissar et al., 2001; Luo and Poeppel, 2007). Indeed, Ahissar and colleagues (2001) found that the synchronization between the speech signal envelope and the MEG signal, presumably originating from the auditory cortex, strongly correlated with the average level of speech comprehension both for normal and compressed speech. They hypothesized that speech envelope synchronization reflected syllable segmentation during sentence presentation and was therefore interpreted as a prerequisite for speech understanding. Moreover, this synchronization with the speech signal in the delta band (1-4 Hz) can predict speech recognition scores at the individual level (Ding et al., 2014). However, it remains controversial whether speech comprehension and envelope tracking are causally linked (e.g., see Zoefel et al., 2018, Kösem and Van Wassenhove, 2017). In a recent study, Ortiz-Barajas and colleagues (2021) have found that newborns possess the neural capacity to track the amplitude and the phase of the speech envelope in their native language (French), as well as in rhythmically similar and different unfamiliar languages (Spanish and English). These findings reveal that envelope tracking represents a basic ability of the auditory system that does not require extensive experience with speech, or knowledge of a given language (i.e., its grammar, or lexicon). These results support the hypothesis that speech envelope tracking may be a necessary prerequisite, although not sufficient, for speech comprehension (Kösem et al., 2016).

In animals, the synchronization of auditory cortex responses with the temporal envelope of guinea pig vocalizations have been observed in several studies (Wallace et al., 2005; Wallace & Palmer, 2009; Grimsley et al., 2011, 2012), which sometimes suggested that cortical responses could be isomorphic to the vocalization envelope (Figure 2A in Grimsley et al., 2012). More recently, Abrams and colleagues (2017) recorded responses of primary auditory cortex neurons at presentation of different levels of speech intelligibility (clear, conversational and compressed) in guinea pigs. They showed that populations of A1 recordings encode both the periodicity and the broadband envelope of the speech signal. These temporal representations in auditory cortex were quite resistant to the degraded conditions (conversational and compressed speech). At the subcortical level, several studies revealed, both in mammals and in birds, that the average responses of inferior colliculus neurons can reflect the communication sound envelope (Suta et al., 2003; Woolley et al., 2006; Rode et al., 2013).

Our main goal was to determine whether the similarities between acoustic envelopes or the loss in envelope tracking ability by auditory neurons, reduce or even prevent the neuronal and behavioral discrimination in situations of acoustic degradations. In a condition-independent scenario, the neurons keep the same ability to track the stimuli envelopes whatever the acoustic conditions (in quiet and in degraded conditions): As long as the stimuli envelopes differ, the neurons will discriminate the stimuli. In contrast, in a condition-dependent scenario, the acoustic degradations reduce the neurons ‘ability to track the stimuli envelopes. In this case, the intense synaptic activity occurring in the auditory nerve when the noise level increases should prevent the central auditory neurons from tracking the stimuli envelopes. To disentangle between these two scenarios, we evaluated the relationship between the envelope tracking of sounds and the neuronal discrimination in the entire auditory system. We simulated auditory nerve fiber (sANF) responses and recorded the neuronal activity in five auditory structures at presentation to four conspecific vocalizations presented in quiet, using three tone-vocoders and two types of noise (a stationary and chorus noise) in anesthetized guinea pigs. We found that subcortical and cortical neurons track the envelopes in the low AM ranges (<20Hz), with relatively high degree of fidelity in original and degraded conditions, suggesting that the auditory system maintains a robust temporal representation from the auditory nerve to the auditory cortex. Behaving mice were also able to discriminate between these communication sounds, and even performed the task above chance level in all noisy conditions. Our results demonstrate that the between-stimulus envelope similarity can explain the changes both in neuronal discrimination at all levels of the auditory system and in behavioral performance in noisy conditions.

## Results

We simulated auditory nerve fiber (sANF) responses and collected neuronal recordings from five auditory structures: the cochlear nucleus (CN), the central nucleus of the inferior colliculus (CNIC), the ventral part of the medial geniculate (MGv), the primary auditory cortex (A1) and a secondary auditory area (VRB).

All analyses were performed on a set of recordings (or simulated recordings) selected using stringent criteria. Note that all the R values presented below are considered as significant (see Method section for more details).

Figure 1A illustrates the overall envelopes of all stimuli in the original, vocoded and noisy conditions. In the following, the term stimulus refers either to the four original or vocoded whistles, or to the four whistles embedded in noise. The four overall envelopes of the stimuli were clearly different between each other in the original and vocoded conditions, however, they progressively became more similar in noisy conditions as the SNR decreased, especially in stationary noise.

**Figure 1.**
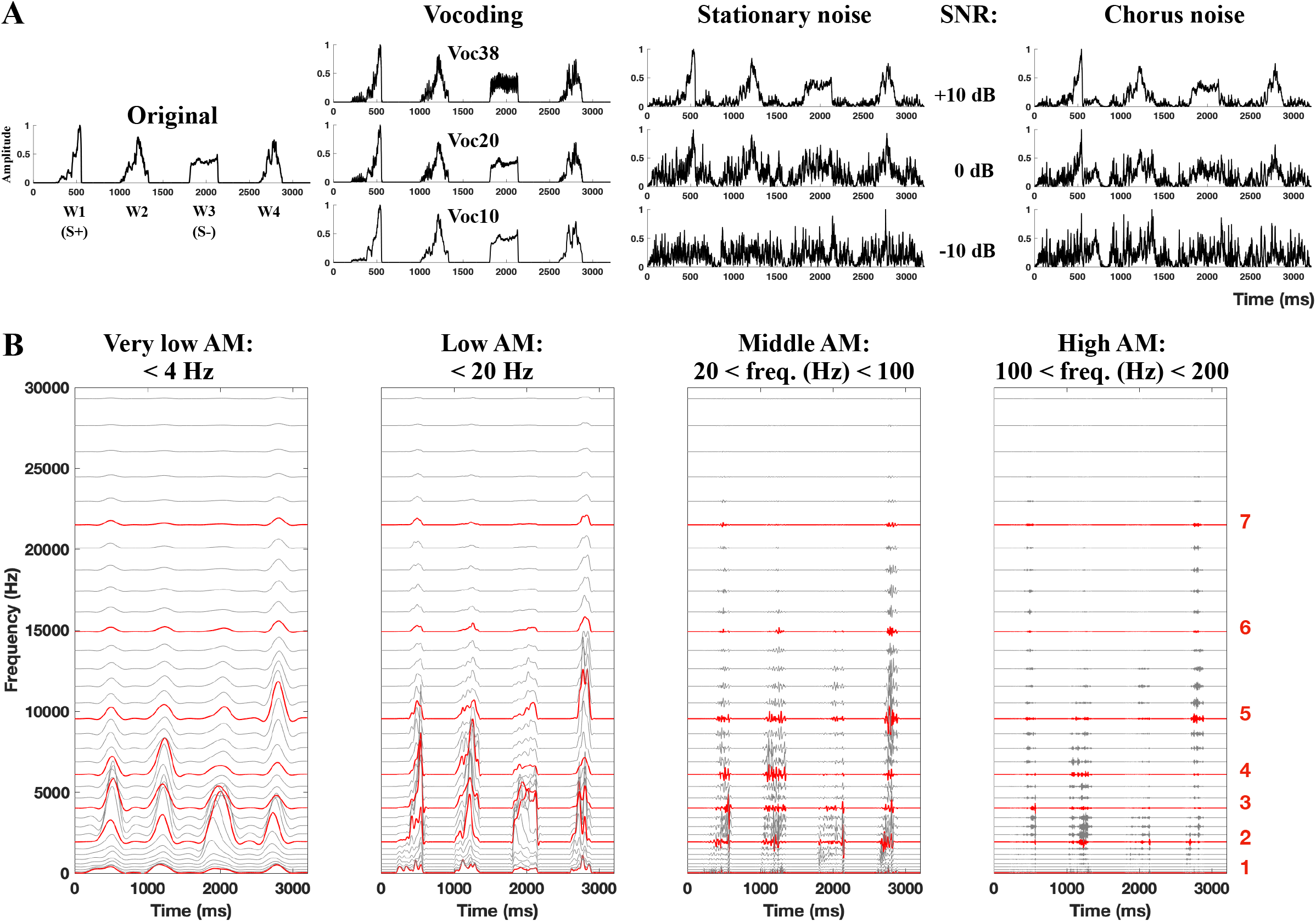
Overall and filtered envelopes in four amplitude modulation ranges. **A.** Overall envelopes of original and degraded stimuli. The envelopes of the four original whistles are presented on the left panel. Two whistles were used for a Go/No-Go behavioral discrimination task (see Fig. 9): the whistle 1 as the “Go or S+” stimulus and the whistle 3 as the “No-Go or S-” stimulus. From left to right, the four envelopes of these stimuli are presented first, in the vocoding conditions (with 38, 20 and 10 frequency bands from top to bottom), then in stationary noise (at +10, 0 and −10 dB SNR from top to bottom) and in chorus noise conditions (at +10, 0 and −10 dB SNR from top to bottom). **B.** Examples of the filtered envelopes for the original vocalizations using a bank of 35 gammatone filters with center frequencies uniformly spaced along a guinea pig - adapted ERB (equivalent rectangular bandwidth) scale ranging from 20 to 30 000 Hz. Four ranges of amplitude modulation (AM) have been investigated here: the very low (<4 Hz), low (<20 Hz), middle (between 20 and 100 Hz) and high (between 100 and 200 Hz) AM ranges. The red curves indicate the seven gammatones selected along the signal for the subsequent analyses.

### Auditory neurons better track the envelopes in the very low and low AM ranges than in fast AM ranges

We first determined which ranges of amplitude modulations are tracked by the subcortical and cortical neurons. To address this question, we filtered the envelopes in four AM ranges: the very low (< 4 Hz), low (< 20 Hz), middle (between 20 and 100 Hz) and high (between 100 and 200 Hz) AM ranges. Figure 1B presents the seven selected gammatones (among the 35) and brings out that the very low and low AM ranges contained larger envelope fluctuations than the middle and high AM ranges. Figures 2A-B present individual examples (Fig. 2A) and populations (Fig. 2B) of PSTHs obtained in each structure at the presentation of the original vocalizations. Based on these PSTHs, it clearly appears that the duration of evoked responses was longer in subcortical structures than in the two cortical areas (A1 and VRB).

**Figure 2.**
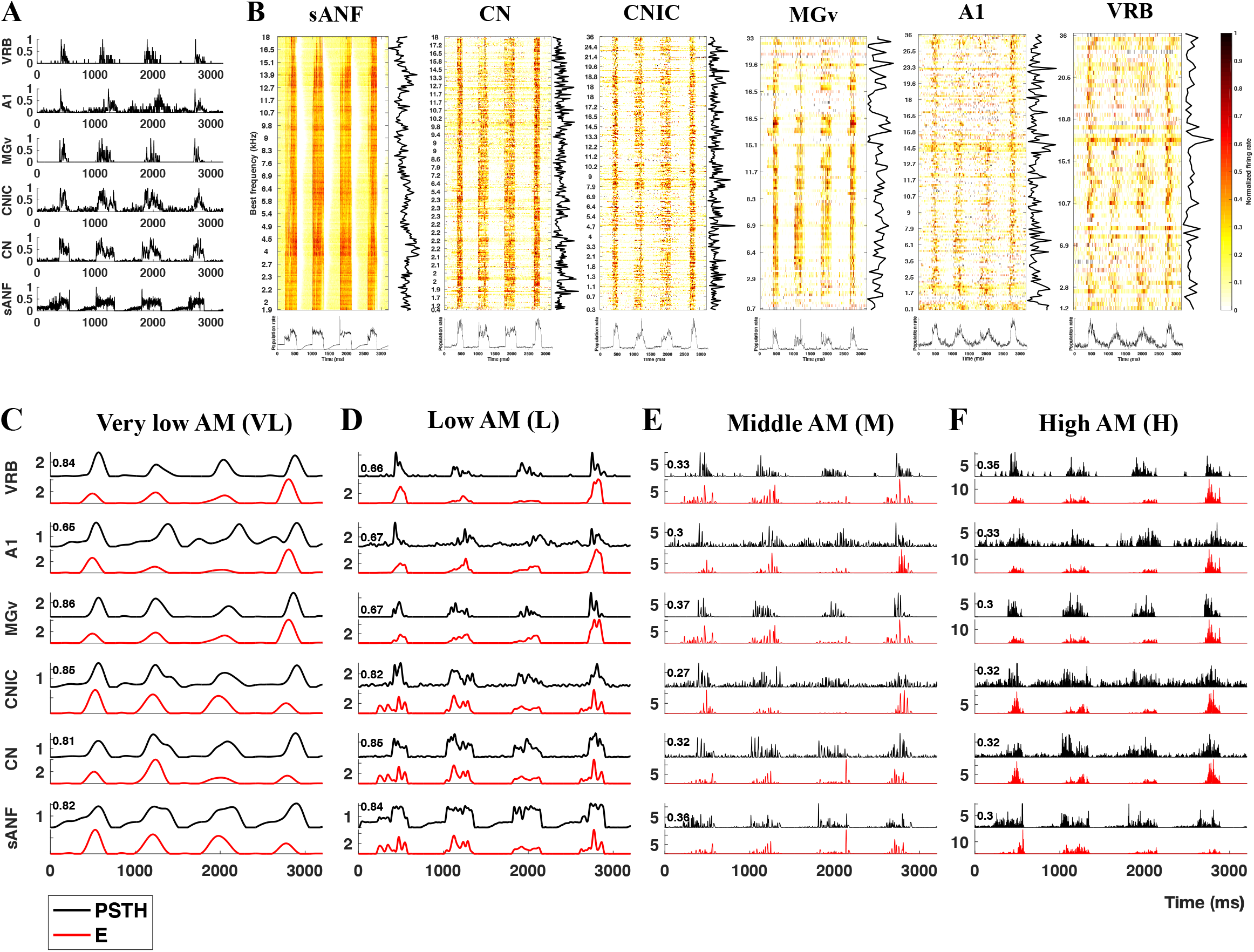
Correlations in the original condition between peri-stimulus time histograms (PSTHs) of subcortical and cortical recordings and the stimulus envelope both filtered in the four selected AM ranges. **A.** Individual examples of the original PSTHs obtained in each structure (from bottom to top, sANF: simulated auditory nerve fibers, CN: cochlear nucleus, CNIC: central nucleus of the inferior colliculus, MGv: ventral division of the medial geniculate, A1: primary auditory cortex, VRB: ventro-rostral belt). **B.** Population responses ranked from the lowest to the highest best frequencies with the color code representing the normalized firing rate. On the bottom of each panel, the population firing rate represents the instantaneous summed activity of the whole virtual population, and on the right, the total firing rate along the different best frequencies. **C-F.** Cross-correlation between the PSTH (in black) and the envelope (in red). In each panel, the PSTHs and the stimulus envelope are filtered in the same frequency range. For each recording, the correlation value between the PSTH and the envelope is shown on the top left. In a given AM range, the stimuli envelopes differ between examples because we selected the gammatone envelope (out of seven gammatones) which induced the highest correlation. Note that the PSTHs are not lagged compared to the envelopes as during the analysis.

Figures 2C-F show the PSTHs from individual recordings (in black) and stimulus envelopes (E, in red) obtained in each structure (and for the sANF) in the original condition, both filtered in the same AM ranges. For each example of E-PSTH, we indicated the extracted cross-correlation value (R) on the top left of each panel. In a given AM range, we represented only the gammatone envelope (out of seven gammatones) which induced the highest correlation. Whatever the structure, in these individual recordings, the higher R values were in the very low and low AM ranges compared with the AM ranges above 20 Hz (middle and high ranges, Fig. 2C-F). The mean R values obtained for each gammatone, each AM range and each structure are presented in Figure 3A. For each central auditory structure, the seven gammatones provided equivalent R values suggesting that auditory neurons did not better track the stimuli’s envelopes in some frequencies than in others. In sANF, gammatone envelopes in lower frequencies (G1, G2) were slightly better in tracking the envelope than those in higher frequencies (Fig. 3A). In the following results, the correlation value selected for each recording at a given AM range was the maximum over the seven gammatones (Rmax_E-PSTH_).

**Figure 3.**
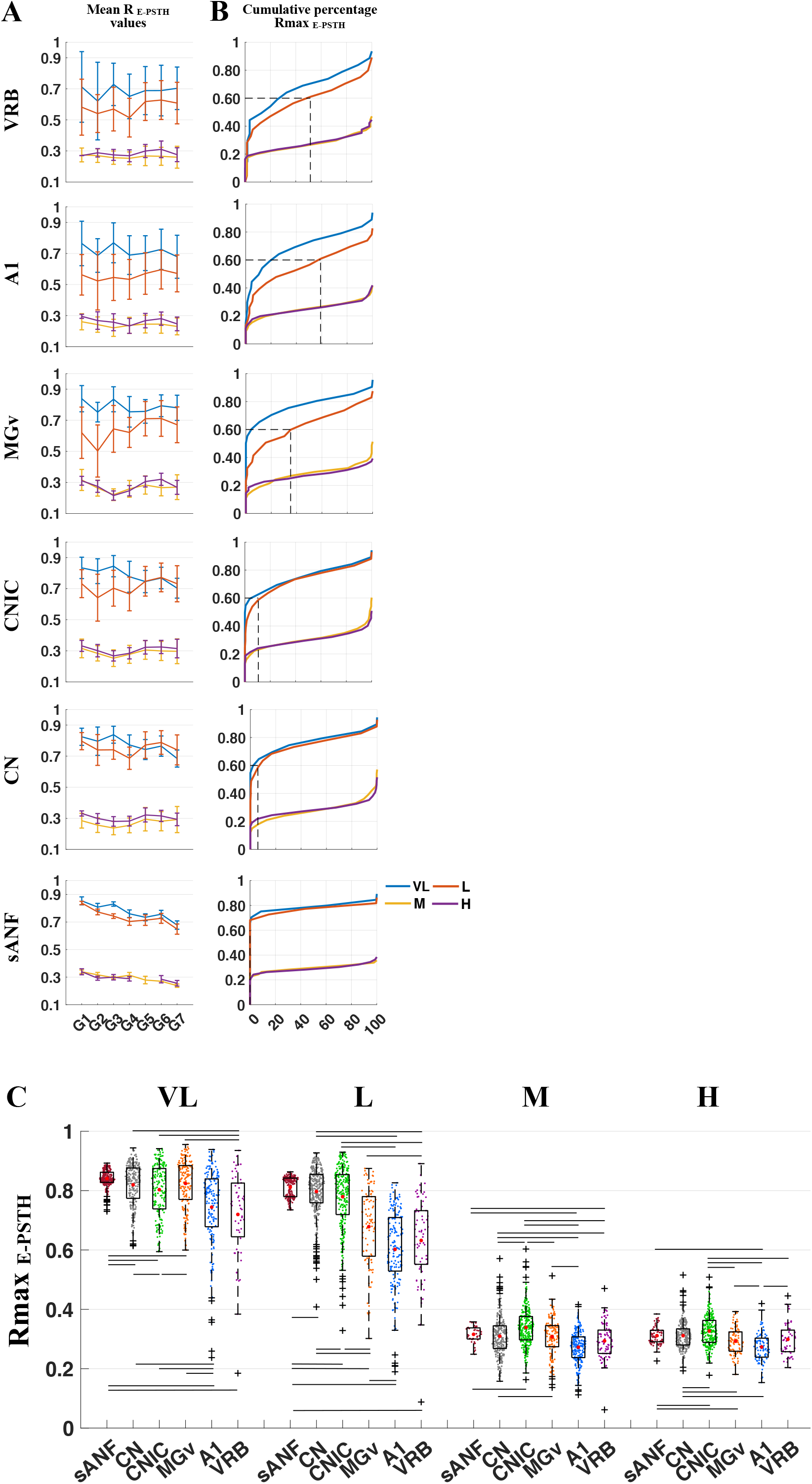
In the original condition, the correlations between the PSTHs and the stimulus envelopes are larger in the very low and low AM ranges in subcortical and cortical structures. **A.** Mean (±STD) correlation values (R_E-PSTH_) in each structure (sANF to VRB), for each AM range (VL, L, M and H, color coded) and for the seven selected gammatones displayed in red in figure 1. **B.** Cumulative distributions of Rmax_E-PSTH_ values for the four AM ranges (color coded) in the six structures (from bottom to top, sANF to cortical areas). The black dotted lines indicated the number of recordings (in percentage) above the R value of 0.6 for the low AM range. **C.** Box plots showing the distributions of the Rmax_E-PSTH_ values for the six auditory structures (sANF to VRB) in the four AM ranges. The red dots correspond to the mean Rmax_E-PSTH_ values. Note the higher Rmax_E-PSTH_ values in the very low (VL) and the low (L) AM ranges compared with the middle (M) and high (H) AM ranges. The black lines represent significant differences between the mean Rmax_E-PSTH_ values (one-way ANOVAs, p<0.05; with post-hoc unpaired t test, lowest p value<0.001).

The cumulative percentages of Rmax_E-PSTH_ values (Fig. 3B) confirmed that the very low and low AM ranges allowed obtaining higher Rmax_E-PSTH_ values than the middle and high AM ranges. Also, it appeared that around 100%, 95%, 90% and 65% of the nerve fibers, cochlear, collicular and thalamic neurons respectively had Rmax_E-PSTH_ values above 0.6 in the low AM range, whereas this was the case for only 40-50% of the cortical neurons (see the dashed lines in Figure 3B).

Figure 3C presents the distributions and the mean Rmax_E-PSTH_ values for each structure in the four AM ranges (VL, L, M and H) in the original condition. Overall, we found a statistically-significant difference in average Rmax_E-PSTH_ values by both the four AM ranges and the six structures (two-way ANOVA, p < 0.05) with a significant interaction between these two factors. For all structures, the mean Rmax_E-PSTH_ values were much higher in the VL and L ranges compared to the M and H ranges. In the VL range, the means of Rmax_E-PSTH_ values in the sANF and three subcortical structures were significantly higher than in the two cortical areas (one-way ANOVA, p < 0.05 with unpaired t test, lowest p value<0.001). In the low AM range, sANF, CN and CNIC recordings displayed significantly higher mean Rmax_E-PSTH_ values than the MGv and cortical recordings (one-way ANOVA, p < 0.05 with unpaired t test, lowest p value<0.001). Similarly, in the middle and high AM ranges, the differences between the structures were less clear but the CNIC recordings still exhibited slightly higher mean Rmax_E-PSTH_ values compared to the other structures. This result was expected for auditory cortex neurons (as they usually exhibit poor abilities to follow fast AM changes) but not expected for subcortical neurons and for sANF (which can synchronize at higher AM rates when tested with periodic artificial stimuli, review in Joris et al., 2004). This suggests that only a partial encoding of faster AM rates contained in complex sounds is performed by subcortical neurons.

To summarize, in the original condition, the neurons’ PSTHs were more strongly correlated with the stimulus envelope in the very low and low AM ranges than in the middle and high AM ranges, both at the subcortical and cortical levels.

### In the original condition, the neuronal discrimination performance is related to the envelope tracking in the faster and lower AM ranges in subcortical and cortical levels, respectively

Does envelope tracking allow auditory neurons to discriminate the four vocalizations in the original condition? To address this question, we looked for relationship between the neuronal discrimination performance and the neurons’ abilities to follow the stimulus envelope (Figure 4). The distributions and mean values of the neuronal discrimination (quantified by the mutual information, MI) are presented for each structure in Figure 4A. As previously reported (Souffi et al., 2020), subcortical neurons (CN, CNIC and MGv neurons) were better in discriminating the original whistles compared to cortical neurons (A1 and VRB neurons, one-way ANOVA p < 0.001 with unpaired t test, highest p value = 0.001 between the cortex and the other structures; Fig. 4A). The scattergrams presented in Figure 4B display the Rmax_E-PSTH_ values as a function of the MI values in each structure and AM range.

**Figure 4.**
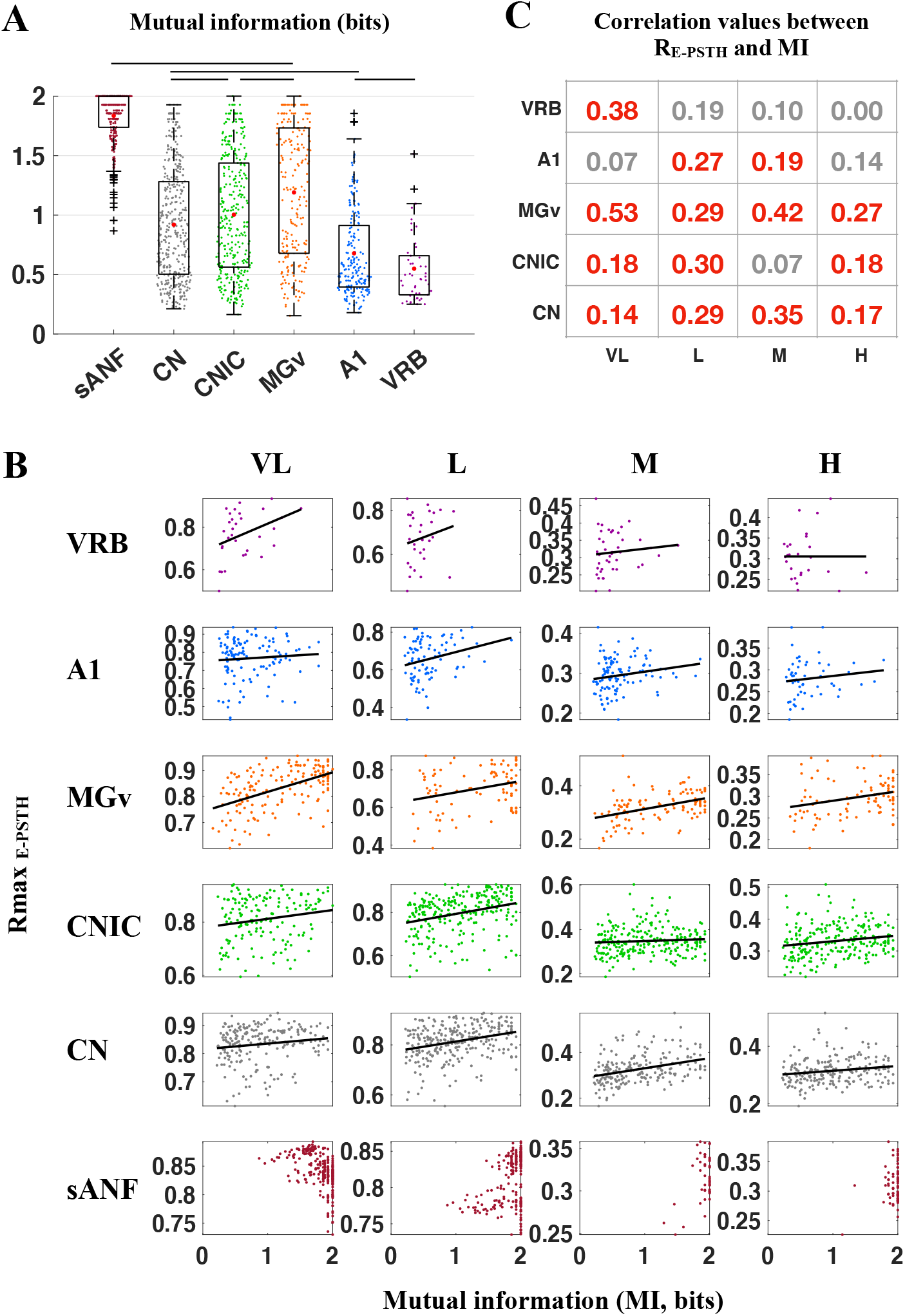
In the original condition, envelope tracking in the very low and low AM ranges better correlates with the neuronal discrimination performance at the cortical and subcortical levels, respectively. **A.** Box plots showing the distributions of the MI values obtained in the six levels of the auditory system in the original condition. The red dots correspond to the mean MI values. Note the lower values obtained at the cortical level in AI and VRB compared to those of the sANF and subcortical structures. The black lines represent significant differences between the mean values (one-way ANOVAs, p<0.05; with post hoc paired t tests, p<0.05). **B.** Scattergrams showing the relationships between Rmax_E-PSTH_ and MI values for the six structures in the four AM ranges. **C.** Matrix summarizing the correlations values between Rmax_E-PSTH_ and MI in each structure and AM range. The values in red indicate that the correlation is significant (p<0.05).

Figure 4C summarizes the correlation values between Rmax_E-PSTH_ and MI in each structure and each AM range. All the significant values of correlation between these two variables are reported in red. In general, significant positive correlations between Rmax_E-PSTH_ and MI values were obtained in all AM ranges in subcortical structures (except in CNIC in the middle AM range) whereas in the two cortical areas, fewer significant correlations were found in the middle and high AM ranges. For the sANF, the range of MI values was too limited to compute reliable correlations (most MI values were close to the maximum of 2 bits). At the subcortical level, the highest correlation values between Rmax_E-PSTH_ and MI values as a whole, were found in MGv. At the cortical level, significant correlations between Rmax_E-PSTH_ and MI values were detected in the low and middle AM ranges in A1 and only in the very low AM range in VRB. Interestingly, each structure presents at least one positive and significant correlation value in the low AM ranges (VL or L) suggesting that the ability for tracking the slow envelopes (<20 Hz) better explains the neuronal discrimination in the entire auditory system.

To summarize, it appeared that in the original condition, the better the tracking of the temporal envelope, the better the between-stimuli neuronal discrimination. For cortical neurons, the correlation between the Rmax_E-PSTH_ and MI values was stronger in the lower AM ranges, whereas for subcortical neurons, there were still significant correlations in the higher AM ranges.

### The acoustic degradations decreased the neuronal discrimination performance but did not affect the envelope tracking performed by the auditory neurons

In almost all situations of acoustic degradations, the neurons’ ability to discriminate between the four vocalizations was decreased. Figures 5A-F present the evolution of the mean MI values for the three levels of degradation conditions: the three tone-vocoders (38, 20 and 10 frequency bands) or the three SNR (+10, 0 and −10 dB SNR) in the two types of noise (stationary and chorus noise). In general, there were only modest effects on the MI values in the vocoding and chorus noise conditions. The decrease was significant only for the 10-band vocoded vocalizations in CN, MGv, and A1 (Fig. 5B, D and E; one-way ANOVA, p<0.05), but it was already significant with 10 or 20-band vocoded vocalizations in the sANF and CNIC respectively (Fig. 5A, C, one-way ANOVA, p<0.05) and no significant difference was detected in VRB. In chorus noise, in CN and CNIC, there was no significant decrease in mean MI values (Fig. 5B, C), whereas in sANF and MGv, the mean MI values significantly decreased at +10 dB or 0 dB SNR respectively (Fig. 5A, D). At the cortical level, there was a significant decrease in A1 at 0 dB SNR and no significant change of mean MI values in VRB (Fig. 5E-F).

**Figure 5.**
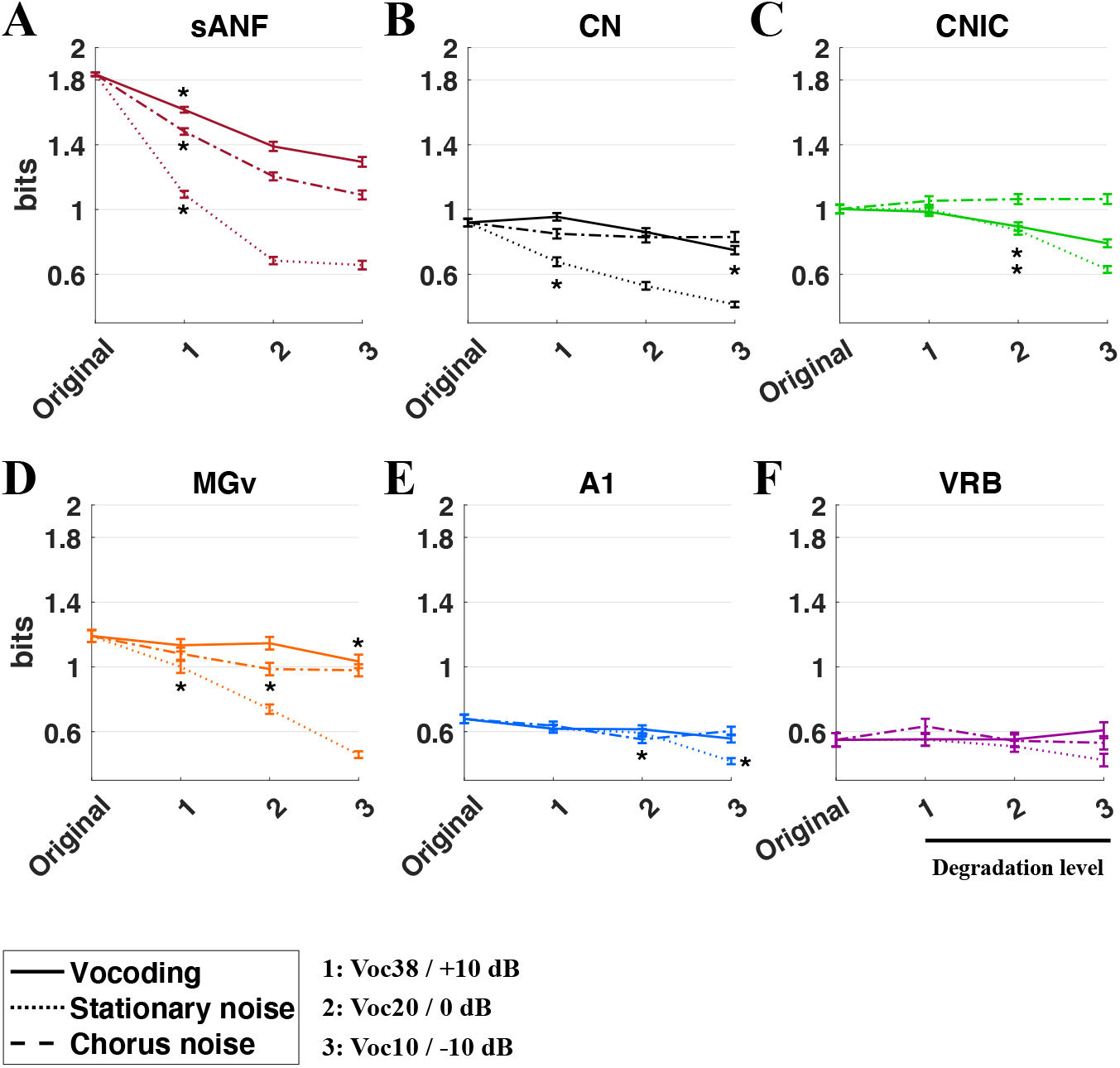
Neuronal discrimination performance in all degraded conditions along the auditory system. **A-F.** Mean neuronal discrimination performance (quantified by the mutual information, in bits) in original condition and in the three situations of acoustic degradation (vocoding in full line, stationary noise in dotted line and chorus noise in dashed line). Note the largest decrease in MI value in the stationary noise in the subcortical structures compared with the relative stability of these values in chorus noise and vocoding. Note also the much smaller decreases observed at the cortical level in the three situations of acoustic alterations. The stars represent significant differences between the mean original values and those obtained in degraded conditions (one-way ANOVAs, p<0.05; with post hoc paired t tests, p<0.05).

Stationary noise strongly reduced the MI values compared to the vocoding and the chorus noise addition. There was a marked difference between the CNIC and the other subcortical structures: the mean MI value in sANF, CN and MGv was significantly reduced already at the +10 dB SNR (Fig. 5A, B, D; p<0.05), whereas the mean MI values in the CNIC was significantly reduced at the 0 dB SNR (one-way ANOVA, p<0.05). At the cortical level, noise significantly reduced the mean MI value in A1 only at the −10 dB SNR (Fig. 5E, one-way ANOVA, p<0.05), whereas the mean MI values in VRB remained unchanged in all conditions (Fig. 5F).

What can be the scenarios explaining the decrease in neuronal discrimination and involving the envelope tracking in situations of acoustic degradations? At least two scenarios can be envisioned: First, in a condition-independent envelope tracking scenario, a neuron keeps the same capacity to track the stimuli envelopes whatever the acoustic conditions, i.e., both in quiet and in conditions of acoustic degradations. In that case, as long as the stimuli envelopes present some differences, the neuron will detect these differences and will discriminate the stimuli. Second, in a condition-dependent scenario, the acoustic degradations reduce the neurons’ ability to track the stimulus envelope. In that case, despite differences between the stimuli envelopes, the intense synaptic activity occurring in the auditory system when increasing the noise level prevents the recorded neuron from tracking the stimuli envelope. To determine which of these two scenarios actually operates, we investigated whether the Rmax_E-PSTH_ values were changed in the conditions of acoustic degradations such as the vocoding or the noise addition (Figures 6–7). Figure 6 shows, for individual recordings, the superpositions of the PSTH and the envelope (E) in the low AM range for the original condition and all the degraded conditions (vocoding, stationary and chorus noise) in each auditory structure. The Rmax_E-PSTH_ values are indicated on the top left of each panel. In these individual recordings, the Rmax_E-PSTH_ values presented very little changes in all degraded conditions compared to the original condition.

**Figure 6.**
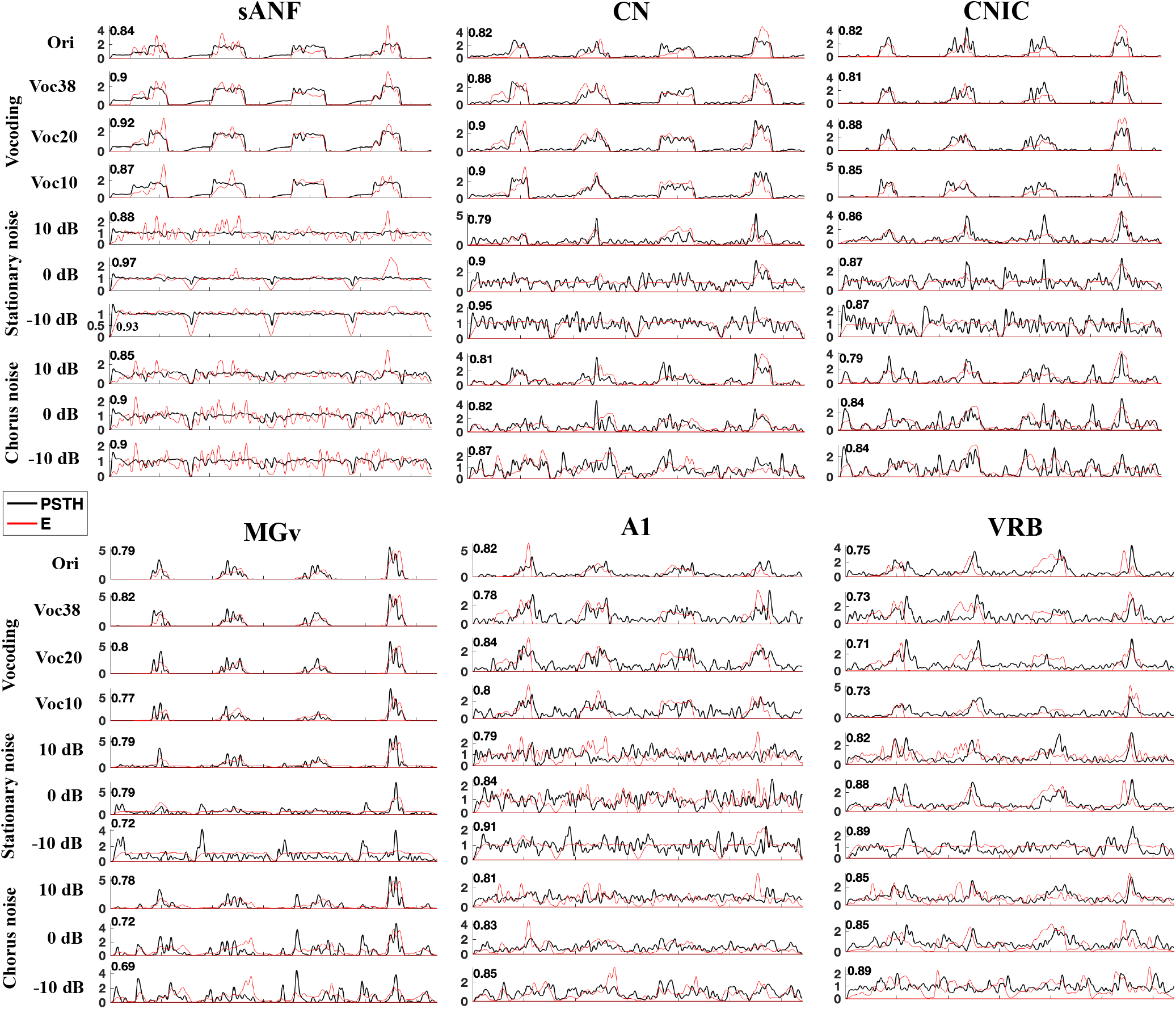
Individual examples of cross-correlation between the PSTH and the envelope in all acoustic conditions and in all structures. Note that for the subcortical and cortical structures, we presented the results from the low AM range. The correlation value between the PSTH (in black) and the envelope (in red) is shown on the left. In all structures, the correlation values for these individual recordings remained similar between the acoustic conditions.

**Figure 7.**
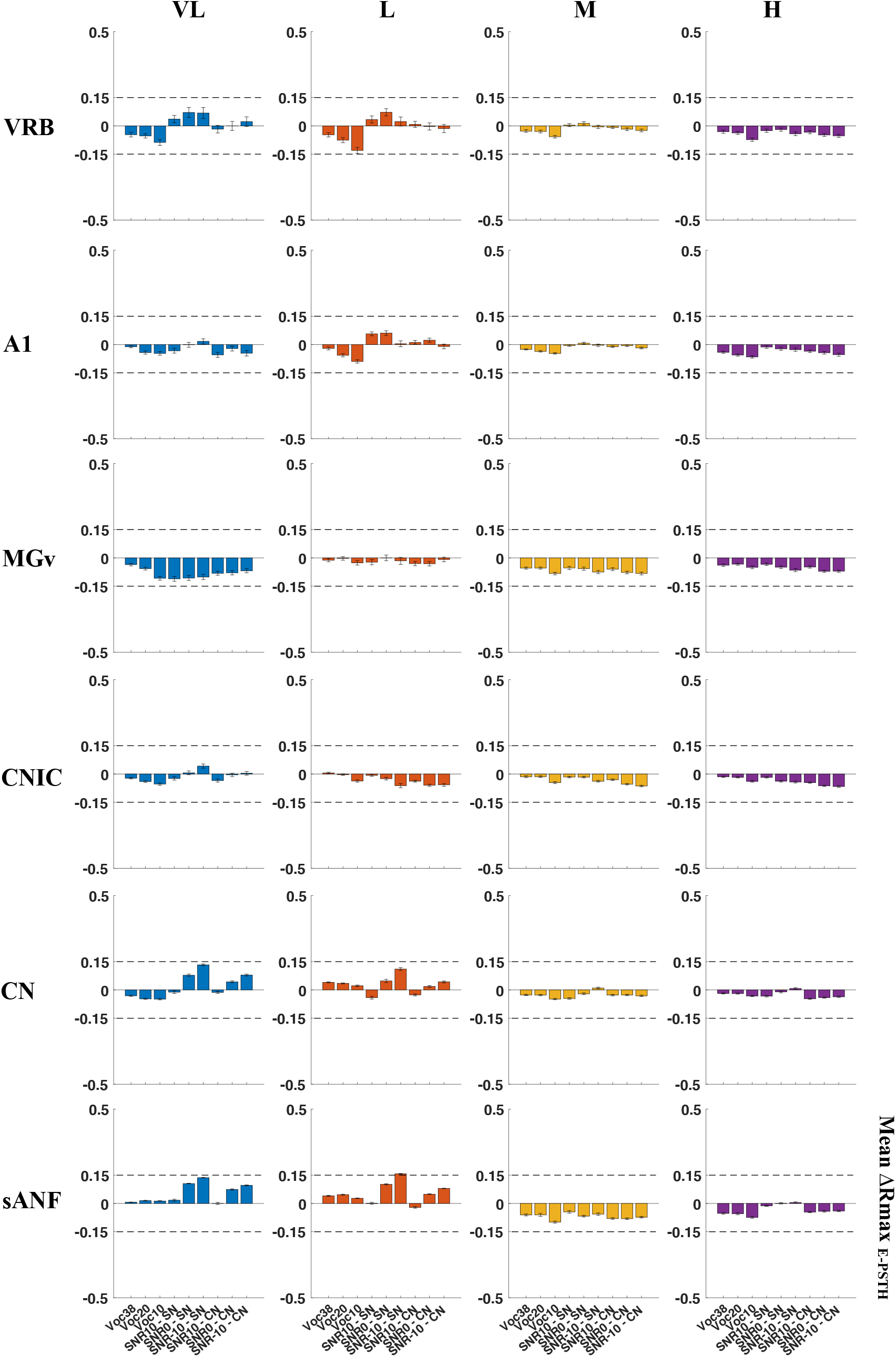
The envelope tracking is only slightly affected by the different situations of acoustic degradation. Mean changes of the Rmax_E-PSTH_ (±SEM) quantified from the original condition in the different situations of acoustic degradations (vocoding, stationary noise (SN) and chorus noise (CN)) for all recordings obtained in the six structures. Note that, in each structure and AM range, the Rmax_E-PSTH_ values were only slightly changed between the original and the degraded conditions.

We next quantified for each recording, the Rmax_E-PSTH_ variations compared with the original condition (ΔRmax). This was quantified in each structure, for all the degraded conditions and each AM range (Fig. 7). Compared with the Rmax_E-PSTH_ values obtained in the original condition, there was little or no change in the degraded conditions for all structures. More precisely, in sANF and CN, we observed a maximal increase in mean Rmax_E-PST_H values of 0.15 and a maximal decrease of 0.007 (and 0.10 for sANF) depending on the degraded conditions and the AM range (Fig. 7). In CNIC and MGv, the mean Rmax_E-PSTH_ changes in degraded conditions were very small (Fig. 7, between −0.06 and 0.04 for CNIC and between −0.11 and −0.002 for MGv). In A1, the changes in Rmax_E-PST_H values varied between −0.09 and 0.06 and in VRB it varied between −0.13 and 0.07.

These results clearly provide evidence that the abilities of neurons for tracking the temporal envelope cues were preserved at each level of the auditory system and in all the situations of acoustic degradations.

### The increase in between-envelope similarity explains the decrease in neuronal discrimination

If the neurons are still able to track the stimulus envelopes in all conditions of acoustic degradations, what could explain the pronounced MI decrease in these situations? The most parsimonious explanation is that the noise addition increases the similarity between stimulus envelopes, which in turns reduces the neuronal discriminative efficiency based on the envelope tracking. We thus quantified the acoustic similarity between the stimulus envelopes in the original condition and in all situations of acoustic degradations. The changes of the between-envelope similarity in each AM range are presented in figure 8A. In the M and H ranges, the between-stimulus envelope similarity was low and remained low in all degraded conditions. In contrast, large changes occurred in the VL and L AM ranges: in these frequency ranges, the envelope similarity increased progressively with the acoustic degradations. In the following results, we will focus on these two AM ranges.

**Figure 8.**
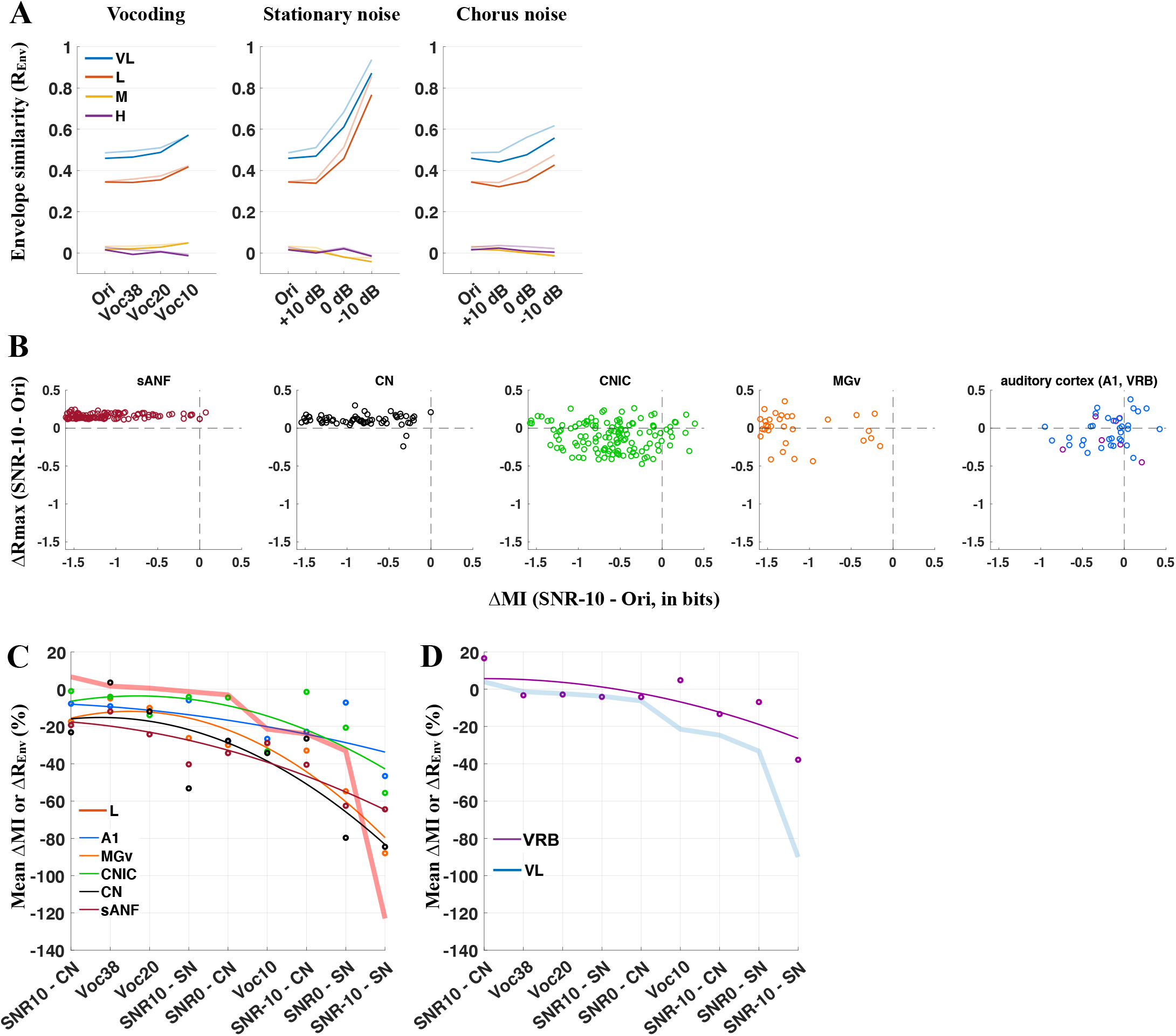
In all situations of acoustic alteration, the decrease in neuronal discrimination performance can be explained by the increase in envelope similarity in the very low and low ranges. **A.** Acoustic similarity (R_Env_) between the envelopes of the four whistles in the original condition (Ori) and in the three situations of acoustic alterations (vocoding, stationary noise and chorus noise) for the very low (VL, blue lines), low (L, red lines), middle (M, yellow lines) and high (H, purple lines) AM ranges. Dark lines correspond to the R_Env_ values based on the 7 selected gammatones, whereas the light lines correspond to the R_Env_ values based on the 35 gammatones. Note that in the stationary noise, the correlation between the stimuli envelopes largely increased in the VL and L ranges, indicating that the stimuli tended to be similar to each others in these AM ranges, which was not the case in the middle and high ranges (M and H). This between-stimuli increase in correlation in the VL and L ranges was much weaker in the vocoding and chorus noise situations. **B.** Scattergrams showing the variation of the maximal correlation (ΔRmax_E-PSTH_) in low AM range as a function of the variation of MI (ΔMI) in −10 dB SNR condition compared to the original condition in each structure. **C.** Mean changes (ΔMI, in percentage) of mutual information in sANF, CN, CNIC MG and A1 and variation (ΔR_Env_, in percentage) of the acoustic similarity in low AM range (red thick line) relative to the original condition. Each dot represents neuronal data (ΔMI) in sANF (in dark red), CN (in black), CNIC (in green), MGv (in orange) and A1 (in blue). From left to right, all degraded acoustic conditions were organized according to the acoustic distance of the envelopes (R_Env_) between the four whistles quantified on Figure 8A. Polynomial fits were generated for the different structures across all degraded conditions (color lines). **D.** Same as in C for the secondary cortical area (VRB) and for the very low AM range (blue thick line).

In the vocoding conditions, the similarity between the four whistle envelopes was relatively constant, except for the 10-band vocoded condition where this similarity was slightly higher. In the stationary noise, the four stimulus envelopes became similar and reached a correlation value of about 0.9 in the −10 dB SNR condition (which is very close to the maximal value of the acoustic similarity). In the chorus noise conditions, the four stimulus envelopes remained different (because spectro-temporal differences were present in the frozen chorus noise) with the highest similarity in the −10 dB SNR condition.

Figure 8B points out that in the condition where the between-stimuli envelope similarity was higher (the −10 dB SNR in stationary noise), the envelope tracking remained similar (the ΔRmax values remained stable) whereas the neuronal discrimination decreased compared to the original condition (most of the ΔMI values are largely negative). This clearly demonstrates the dissociation between changes in Rmax value and those in MI value. Figures 8C-D highlight the close relationship between the acoustic similarity of the four stimulus envelopes and the abilities of auditory neurons to discriminate them. In the subcortical structures and in A1, as the acoustic distance between the four stimulus envelopes in the low AM ranges progressively decreased, the neuronal discrimination decreased (Fig. 8C). Similarly, in the secondary area VRB, as the acoustic distance in the very low AM ranges between the four stimulus envelopes progressively decreased, the neuronal discrimination of cortical neurons decreased (Fig. 8D).

Together, these results indicate that it is not a loss in neuronal envelope tracking which leads to a reduction of the neuronal discriminative abilities in the degraded conditions rather, it is the increase of the envelope similarity in situation of acoustic degradations that is responsible for the decrease in discrimination abilities. Thus, the between-stimuli envelope similarity in the lower AM ranges (<20 Hz) could be used as an “acoustic marker” for predicting the evolution of the discrimination in the entire auditory system.

### The increase in between-envelope similarity also controls the behavioral performance

To determine whether the discrimination performance of auditory neurons provide a neuronal basis for behavioral performance, we tested whether behaving animals can discriminate between whistles when engaged in a operant conditioning task involving the same stimuli. We opted to train mice in a behavioral task rather than guinea pigs for two main reasons: (1) guinea pigs are poor and slow learners in instrumental tasks (2) this avoided that the stimuli used for the behavioral task have particular meanings.

The behavioral task was a Go/No-Go task involving the discrimination between two of the four whistles used in our electrophysiological studies: Licks to the S+ were rewarded by a 0.5μL drop of water and licks to the S-were punished by a 5-second time-out period (Fig. 9A). Mice were first trained for 5-10 initial sessions to perform the discrimination in the original condition until they reached 80% of correct responses for two successive days. Then, the mice were sequentially trained in the stationary noise at the +10, 0 and −10 dB SNR for at least four sessions. The performance at the last four ones at each SNR are displayed on figure 9B. For all mice, the performance decreased at the 0 and −10 dB SNR, even if some mice were still at 80% of correct performance whereas others were at the chance level. Surprisingly, in the chorus noise, the performance of most of the mice were relatively stable, which can be explained by the fact that acoustically the chorus noise surrounding the two target vocalizations differed between the two whistles, so that there were more acoustic cues to discriminate between the target stimuli in these conditions. Despite this pitfall, the main result of this behavioral study was that mice can discriminate the target vocalizations above chance level even at the −10 dB SNR in stationary noise. Furthermore, the decrease in behavioral performance was proportional to the reduction of the differences between the two temporal envelopes in the low AM range (Fig. 9C): when the differences in temporal cues between two whistles dropped by more than 10%, the behavioral performance was impacted (last two points on figure 9C). These results provide evidence that the mice behavioral performance is potentially under the control of temporal envelope cues in the low AM range.

**Figure 9.**
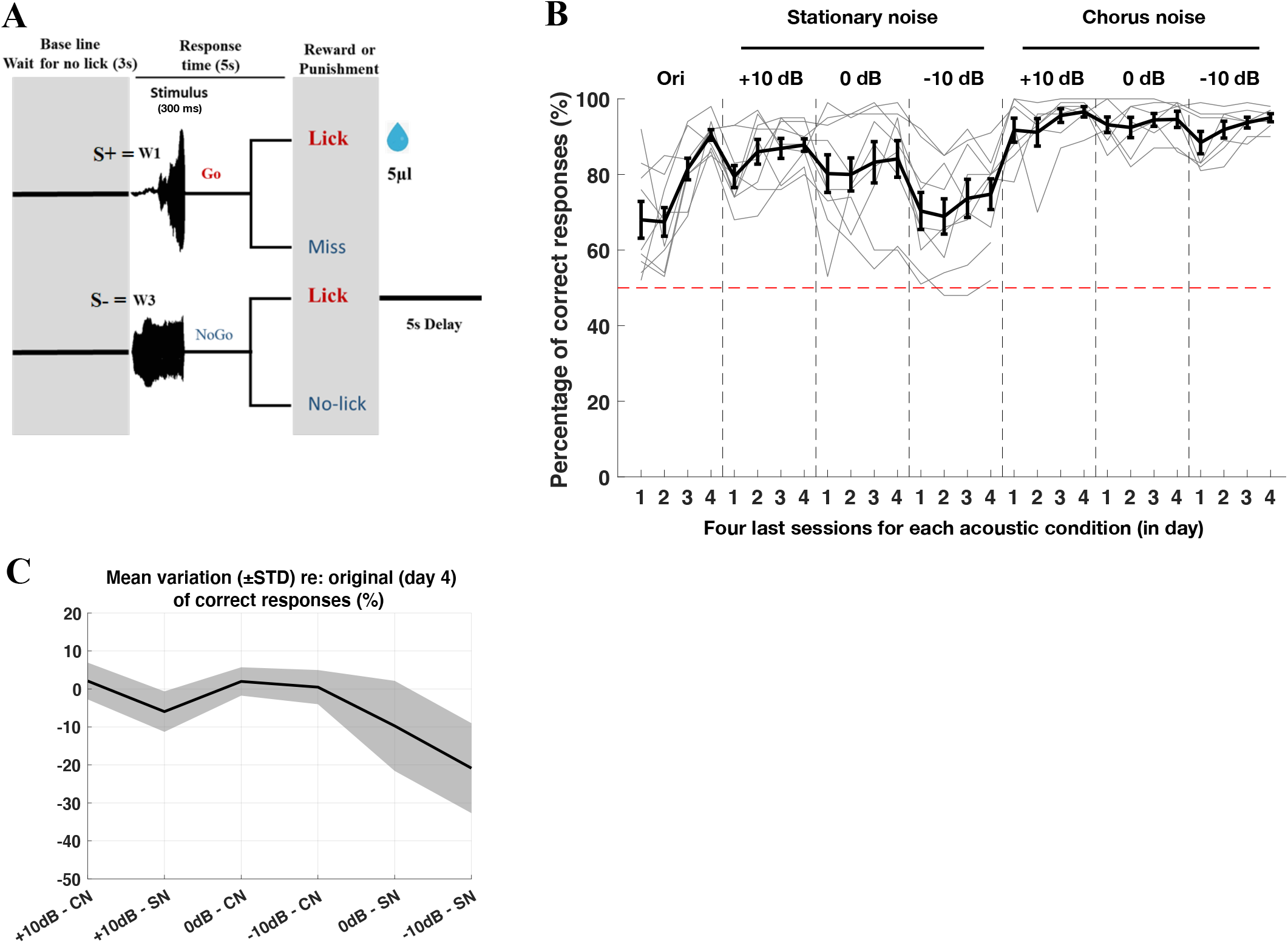
Behavioral discrimination performance in original and noise conditions in mice. **A.** Diagram summarizing the Go/No-Go behavioral task. At the presentation of the S+ signal (Whistle 1), the mice have five seconds to lick for obtaining a drop of water. At the S-signal (Whistle 3), a lick response triggers a five second time-out period. A three second baseline period with no lick is required before the beginning of each trial. **B.** Percentage of correct responses obtained during the four last sessions for each condition. The dark thick line corresponds to the mean (±SEM) values obtained for all mice. The individual performances of each mouse (n=9) are presented by the grey thin lines. The last four sessions of discrimination in the original conditions are represented followed by the discrimination in the three conditions in stationary noise (+10, 0 and −10 dB SNR), followed by the discrimination in the three conditions in chorus noise (+10, 0 and −10 dB SNR). The chance level is represented by the red dashed line. **C.** Mean variation of correct responses compared to the performance obtained for each mice at the last day in the original conditions. All degraded conditions are ranked according to the acoustic similarity in low AM range between the W1 and W3. The only drops in performance were observed for the 0 dB and the −10 dB SNR in the stationary noise.

## Discussion

Our first major result is that the neuronal discrimination performance in the original condition was correlated with the capacity for tracking the envelopes in the low AM range both for subcortical and cortical neurons, except in secondary auditory cortex (VRB) where this close relationship was found only in the very low AM range. Our second major result is that, in each structure, the ability for envelope tracking under acoustic degraded conditions only slightly changed compared to the original condition. Overall, our findings revealed that the decrease in neuronal and behavioral discrimination can be explained by the increased similarity between the stimuli envelopes in the low AM ranges (<20 Hz).

### Slow envelope tracking: a general property of auditory neurons

Previous electrophysiological studies in anesthetized animals have shown that cortical responses can be partially predicted by the temporal envelope of communication sounds (Wang et al., 1995; Bar-Yosef et al., 2002; Nagarajan et al., 2002; Grimsley et al., 2012; Abrams et al., 2017). For example, both in marmosets and guinea pigs, it was reported that A1 neuronal populations phase-locked with the envelope of conspecific vocalizations having a periodic temporal structure (marmoset: Wang et al., 1995; guinea pig: Grimsley et al., 2012). In guinea pig, high correlations were found in A1 and in two non-primary cortical areas (VRB and S), while in marmosets some A1 neurons can even phase-lock despite acoustic modifications (time-compressed, time-expanded, time-reversed).

In the original non-degraded condition, our results are in line with these previous studies. Indeed, we found that A1 and VRB neurons exhibited high correlation values in lower AM ranges (<20 Hz) highlighting that low-pass filtered cortical responses represent the slow amplitude modulation information. Compared to previous studies, we filtered the envelopes and the neuronal responses in the same frequency bands for direct comparisons between neuronal responses and acoustic temporal information. These relationships were investigated in four ranges of amplitude modulations, from very low (< 4 Hz) to high (100 and 200 Hz) ranges. This procedure allowed investigating the properties of the envelope tracking performed by auditory neurons in a more direct way. Furthermore, we compared the degree of envelope tracking performed by subcortical and cortical neurons in challenging situations where the envelope is either relatively well preserved or strongly degraded. In anesthetized marmosets, Nagarajan and colleagues (2002) found that the synchronization between the responses of A1 neurons and the temporal envelope of vocalizations was highly significant and, interestingly, this property was underestimated based on responses to amplitude-modulated tones. In addition, they pointed out that A1 responses were quite resistant to spectral degradations (generated by a noise-vocoder) and to background noise addition up to 0 dB SNR. More importantly, the responses were similar when the vocalization temporal envelope was preserved between 2 and 30 Hz, whereas the responses were strongly reduced when the envelope was low-pass filtered at 4 or 10 Hz (Nagarajan et al., 2002). We confirmed these cortical results on several aspects. First, we detected higher correlation values in the lower AM ranges (<20 Hz) for each tested acoustic condition. Second, we showed that this capacity for envelope tracking was little affected by the presence of noise addition or by the vocoding. Here, we also extended these results to a non-primary cortical area (VRB), to each subcortical level, and even to sANF. Note that this property is not specific to the processing of conspecific vocalizations: similar results were found in the auditory cortex of anesthetized guinea pigs (Abrams et al., 2017) with noisy speech signals (called conversational). The study by Abrams and colleagues showed that A1 neurons represent the envelope (<50 Hz) of speech signals with a high fidelity (R>0.5) in different conditions (clear, conversational and compressed). Together, these results highlight that subcortical and cortical auditory neurons maintain their capacity to track the slow envelope of natural sounds both when they are composed of noise-free vocalizations or a mixture of noise and vocalizations, suggesting that this property is immutable and unchanged by the acoustic degradations.

The subcortical processing of communication sounds has probably long been underestimated. Only a few electrophysiological studies have shown that subcortical neurons can display responses very close to the envelopes of natural stimuli (in the inferior colliculus: Suta et al., 2003; Rode et al., 2013; thalamus: Tanaka and Taniguchi, 1991; Philibert et al., 2005; Suta et al., 2007). Using a non-linear model (Dual-Resonance Non-Linear model) to simulate the peripheral processings and to extract the stimuli envelopes (<100 Hz), Rode and colleagues (2013) found that between 15 and 60% collicular neurons displayed high correlation for at least one of the three envelope vocalizations, and a subset of collicular neurons even followed the envelopes of these three guinea pig vocalizations with high correlation values (>0.85). A similar range of correlations (between 0.6-0.9) in CNIC was obtained in the present study and, as in their study, we also did not find a relation between the preferred gammatone (eliciting the highest R value) and the best frequency of the neurons.

Using artificial stimuli, many studies have shown that subcortical neurons can discharge at higher AM rates compared to cortical neurons (Creutzfeldt et al., 1980; Frisina et al., 1990; Rhode and Greenberg, 1994; Neuert et al., 2001; for review, Joris et al., 2004). In the low AM range (<20 Hz), we noticed a decrease in mean correlation (Rmax_E-PSTH_) values from midbrain to thalamus and to cortex (Fig. 3C) reflecting that the further away from the periphery, the less precise is the phase-locking ability. For higher AM rates, we expected higher correlations between the neuronal responses and the envelopes for subcortical neurons. Surprisingly, such a hierarchy was not detected in our results, the mean correlations in higher AM rates (>20 Hz) being similarly low for each structure including the sANF. One hypothesis is that shorter segments of neuronal responses could be highly correlated to the higher AM ranges of the envelopes. If so, reducing the time window on which the correlation is computed should increase the correlations in the higher AM ranges. We computed the cross-correlation for each whistle (around 300 ms) and still found low correlation values in higher AM ranges (data not shown). This suggests that if higher correlation values exist in higher AM ranges, smaller temporal windows (less than several hundreds of milliseconds) are required to reveal them. The fact that Abrams and colleagues (2017) found some residues of the fundamental frequency (between 100-120 Hz, relative to the pitch) in segments of A1 responses no longer than 100 ms argues in favor of this possibility.

### The decrease in neuronal discrimination can be explained by the increase of between-envelopes similarities in the low AM range

In the original condition, the neuronal discrimination performance was significantly correlated with the envelope tracking in the low AM range (<20 Hz) for all structures, except in VRB where this relation was significant in the very low AM range (<4 Hz). This indicates that, in all structures, the more neurons track the slow envelope (<20 Hz), the higher the neuronal discrimination performance. Given that the neurons still followed the stimulus envelope in the degraded conditions, it was crucial to determine if the modifications of the four envelopes in each AM range predict the decrease of the neuronal discrimination in these conditions.

In situations of acoustic degradation, the envelopes of the original stimuli were strongly altered leading to situations where the envelopes were mostly dominated by the noise envelopes. However, the three situations of acoustic degradations used here notably differed. In the tone-vocoder situation, the vocoding strongly degraded the spectral content but preserved the slow temporal envelope (Shannon et al., 1995; Kates 2011; Souffi et al., 2020) and thus the main temporal differences between the four vocalizations are preserved. In the chorus noise, there was only a small increase in acoustic similarity in the low AM range because the chorus noise itself contains strong temporal variations which differ from one whistle to another. Therefore, when the target vocalizations were inserted in the chorus noise, specific regions in the spectro-temporal domain were dominated by the target vocalizations, while in other regions, it was dominated by the chorus noise. Consequently, the target vocalizations embedded in the chorus noise generate stimuli that can be discriminated at all SNRs either based on the vocalization envelopes or based on the chorus noise envelope itself. In all structures, the neuronal discrimination showed little decrease in the chorus noise (see Fig. 5) and so was the behavioral performance (Fig. 9B). It was only in the stationary noise that the four envelopes became closer as the level of degradation increased, and this was true only in the lower AM ranges (< 4 Hz and < 20 Hz, see Fig. 8A). This was detrimental for discriminating the degraded stimuli in noisy conditions. In such situations, envelope tracking become inefficient and worst, can strongly reduce the neuronal discrimination along the auditory system. Therefore, reducing or increasing the envelope differences in the low AM range would constrain or facilitate the neuronal discrimination in subcortical and cortical levels. Furthermore, the behavioral performance of mice for discriminating two of the guinea pig vocalizations revealed that they can discriminate the target vocalizations in quiet (with > 90% correct performance) and can even discriminate the vocalizations up to 0 dB SNR in stationary noise (with 70-80% correct performance), suggesting that the between-stimuli envelope differences potentially explains the behavioral performance during a discrimination task. Previous studies have reported good behavioral discrimination performance in conditions of acoustic degradations such as vocoded consonants or vowels (Ranasinghe et al., 2012a, b), consonant sounds in various levels of background noise (Shetake et al., 2011), conspecific bird songs embedded in stationary and chorus noise (Narayan et al., 2007) or in broadband dynamic moving ripples (Homma et al., 2020). In all these studies, the discrimination performance of auditory cortex neurons, based upon spike-timing, has been found to match relatively well the behavioral performance (Narayan et al., 2007; Ranasinghe et al., 2012a; Homma et al., 2020) and sometimes even with performance of human subjects (Walker et al., 2008). Our present results extend this relationship in several aspects. First, we found that the decrease in neuronal discrimination can be explained by the between-stimuli envelope differences in the low AM range (< 20 Hz). Indeed, we showed that whatever the initial level of neuronal discrimination - which was higher for subcortical than for cortical neurons - increasing the between-stimuli envelope similarity impacted the neural performance at each level of the auditory system. This indicates that it is not a loss in neuronal envelope tracking which leads to a reduction of the neuronal discriminative abilities in the degraded conditions, rather, it is the direct consequence of the acoustic distance changes between stimulus envelopes in the degraded conditions. Last, when these acoustic changes reach a certain level (about 20% or more), this impacted on the behavioral discrimination performance. Together, these results emphasize the crucial role of AM cues in the neural and behavioral performance.

### Comparison with human studies: case of newborn infants

In humans, it has been claimed that the tracking of the speech envelope plays a key role in speech comprehension. Speech envelope corresponds to the slow amplitude fluctuations of the signal over time, with peaks occurring roughly at the syllabic rate. The two pioneer results supporting this view are (1) that comprehension is impaired when the speech envelope is filtered out (Drullman et al., 1994a, b), and (2) that adult listeners readily understand degraded speech in which only the envelope is preserved, at least when speech is presented in silence (Shannon et al., 1995). Additionally, studies have shown that when adults listen to speech, their neuronal activity synchronize with specific features of the speech envelope, a phenomenon known as speech envelope tracking (Ahissar et al., 2001; Luo and Poeppel, 2007; Abrams et al., 2008; Nourski et al., 2009). One feature with which brain activity may synchronize is the amplitude of the speech signal: amplitude synchronization occurs when the amplitude of the neural activity in the auditory cortex follows the contours of the speech envelope (Abrams et al., 2008; Nourski et al., 2009; Kubanek et al., 2013). The auditory cortex also synchronizes with the phase of the speech envelope by modulating the phase of its ongoing oscillations to match the phase of the envelope (i.e., phase-locking) (Peelle et al., 2013; Pefkou et al., 2017). The quality of amplitude (Ahissar et al., 2001; Nourski et al., 2009) and phase synchronization (Luo and Poeppel, 2007; Peelle et al., 2013) has been found to correlate with speech comprehension. These findings led to the view that speech envelope tracking is a key mechanism in speech comprehension.

Several recent electrophysiological results have provided new insights into this putative speech envelope tracking mechanism. First, oscillations whose frequency corresponds to the modulation frequency of the speech envelope (4-5 Hz) has been found to be independent of comprehension: brain responses in the theta band track the speech envelope even when speech is time-compressed at a rate that renders it incomprehensible for adult listeners (Pefkou et al., 2017; Zoefel and VanRullen, 2016; Kösem et al., 2016). Results from newborns and young infants have also brought new insights in this field. For example, combining hemodynamic (near-infrared spectroscopy) and EEG recordings to measure brain responses to syllables differing in consonants, Cabrera and Gervain (2020) showed that infants (9-10 months old) detect consonant changes on the basis of envelope cues (without the temporal fine structure) and they can even do so on the basis of the slow temporal variation alone (AM <8 Hz). These results are consistent with behavioral data obtained with older infants and adults for whom the slowest envelope cues are also sufficient to detect consonant changes in silence (Drullman, 1995; Shannon et al., 1995; Cabrera et al., 2015; Cabrera & Werner, 2017). More recently, by recording EEG from French newborns (within their five first days of life) during presentation of sentences in English, French and Spanish having the same amplitude and frequency modulation spectra in the three languages, it was shown that speech envelope tracking occurs at birth (Ortiz-Barajas et al., 2021). This suggests that the cortical networks of newborns (exclusively exposed to French before birth) have the capacity to track the amplitude and the phase of speech envelope in their native language as well as in unfamiliar language (Spanish and English). Altogether, these results suggest that amplitude- and phase-tracking take place in the absence of attention and comprehension.

Thus, envelope tracking can be viewed as a universal mechanism used in all species to discriminate between communication sounds in a large diversity of acoustic situations ranging from quiet to adverse, challenging, conditions.

## Limitations of the study

Obviously, one of the main limitations of the present results is that the neural results were not obtained on the same animals as those performing the behavioral task. So, we cannot know what is the link between the changes in neural discrimination performance occurring in a given level in the auditory system and the behavioral performance. Very recently, it was reported that different situations of acoustic degradation (similar to ours) trigger different neuronal mechanisms when human subjects had to discriminate between word and non-word stimuli (Hernandez-Perez et al., 2021). The discrimination between vocoded stimuli significantly activated the medial olivocochlear (MOC) reflex, proportionally to task difficulty, potentially via corticofugal connections. By contrast, in speech-shaped noise and babble noise (equivalent to our stationary and chorus noise) the discrimination between target stimuli did not activate the MOC reflex but increased both the global midbrain and cortical responses. Thus, it is possible that during a challenging discrimination task different neural mechanisms are operating to discriminate communication sounds depending on the degraded conditions. These human findings, obtained with global responses, urge for more detailed studies to better understand the neural mechanisms involved in controlling our perceptive abilities in challenging conditions.

## Materials and Methods

Most of the Methods are similar to those described in Souffi and colleagues (2020).

### Subjects

These experiments were performed under the national license A-91-557 (project 2014-25, authorization 05202.02) and using the procedures N° 32-2011 and 34-2012 validated by the Ethic committee N°59 (CEEA, (Comité d’Ethique pour l’Expérimentation Animale) Paris Centre et Sud). All surgical procedures were performed in accordance with the guidelines established by the European Communities Council Directive (2010/63/EU Council Directive Decree).

Extracellular recordings were obtained from 47 adult pigmented guinea pigs (aged 3 to 16 months old, 36 males, 11 females) at five different levels of the auditory system: the cochlear nucleus (CN), the inferior colliculus (IC), the medial geniculate body (MGB), the primary (A1) and secondary (area VRB) auditory cortex. Animals, weighting from 515 to 1100 g (mean 856 g), came from our own colony housed in a humidity (50-55%) and temperature (22-24°C)-controlled facility on a 12 h/12 h light/dark cycle (light on at 7:30 A.M.) with free access to food and water.

Two days before the experiment, the animal’s pure-tone audiogram was determined by testing auditory brainstem responses (ABR) under isoflurane anesthesia (2.5%) as described in Gourévitch and colleagues (2009). A software (RTLab, Echodia, Clermont-Ferrand, France) allowed averaging 500 responses during the presentation of nine pure-tone frequencies (between 0.5 and 32 kHz) delivered by a speaker (Knowles Electronics) placed in the animal right ear canal. The auditory threshold of each ABR was the lowest intensity where a small ABR wave can still be detected (usually wave III). For each frequency, the threshold was determined by gradually decreasing the sound intensity (from 80 dB down to −10 dB SPL). All animals used in this study had normal pure-tone audiograms (Gourévitch et al., 2009; Gourévitch and Edeline, 2011).

### Acoustic stimuli

The acoustic stimuli were the same as in Souffi and colleagues (2020, 2021). They were generated using MatLab, transferred to a RP2.1-based sound delivery system (TDT) and sent to a Fostex speaker (FE87E). The speaker was placed at 2 cm from the guinea pig’s right ear, a distance at which the speaker produced a flat spectrum (± 3 dB) between 140 Hz and 36 kHz. Calibration of the speaker was made using noise and pure tones recorded by a Bruel and Kjaer microphone 4133 coupled to a preamplifier BandK 2169 and a digital recorder Marantz PMD671.

The Time-Frequency Response Profiles (TFRP) were determined using 129 pure-tone frequencies covering eight octaves (0.14-36 kHz) and presented at 75 dB SPL. The tones had a gamma envelop given by 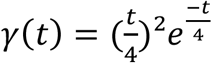, where *t* is time in ms. At a given level, each frequency was repeated eight times at a rate of 2.35 Hz in pseudorandom order. The duration of these tones over half-peak amplitude was 15 ms and the total duration of the tone was 50 ms, so there was no overlap between tones.

A set of four conspecific vocalizations was used to assess the neuronal responses to communication sounds. These vocalizations were recorded from animals of our colony. Pairs of animals were placed in the acoustic chamber and their vocalizations were recorded by a Bruel & Kjaer microphone 4133 coupled to a preamplifier B&K 2169 and a digital recorder Marantz PMD671. A large set of whistle calls was loaded in the Audition software (Adobe Audition 3) and four representative examples of whistle were selected. As shown in figure 1A (left panels), their overall envelopes clearly differed.

The four selected whistles were processed by three tone vocoders (Gnansia et al., 2009, 2010). In the following figures, the unprocessed whistles will be referred to as the original versions, and the vocoded versions as Voc38 (Voc20, Voc10 respectively) for the 38-band (20-band, 10-band, respectively) vocoded whistles. In contrast to previous studies that used noise-excited vocoders (Nagarajan et al., 2002; Ranasinghe et al., 2012; Ter-Mikaelian et al., 2013), a tone vocoder was used here, because noise vocoders were found to introduce random (i.e., non-informative) intrinsic temporal-envelope fluctuations distorting the crucial spectro-temporal modulation features of communication sounds (Shamma and Lorenzi, 2013; Kates, 2011; Stone et al., 2011).

Figure 1A displays the overall envelopes of the 38-band vocoded (first row, second panel), the 20-band vocoded (second row, second panel) and the 10-band vocoded (third row, second panel) of the four whistles. The three vocoders differed only in terms of the number of frequency bands (i.e., analysis filters) used to decompose the whistles (38, 20 or 10 bands). The 38-band vocoding process is briefly described below, but the same principles apply to the 20-band or the 10-band vocoders. Each digitized signal was passed through a bank of 38 fourth-order Gammatone filters (Patterson, 1987) with center frequencies uniformly spaced along a guinea-pig adapted ERB (Equivalent Rectangular Bandwidth) scale ranging from 50 to 35505 Hz (Sayles and Winter, 2010). In each frequency band, the temporal envelope was extracted using full-wave rectification and low-pass filtering at 64 Hz with a zero-phase, sixth-order Butterworth filter. The resulting envelopes were used to amplitude modulate sine-wave carriers with frequencies at the center frequency of the Gammatone filters, and with random starting phase. Impulse responses were peak-aligned for the envelope (using a group delay of 16 ms) and the acoustic temporal fine structure across frequency channels (Hohmann, 2002). The modulated signals were finally weighted and summed over the 38 frequency bands. The weighting compensated for imperfect superposition of the bands’ impulse responses at the desired group delay. The weights were optimized numerically to achieve a flat frequency response.

The four whistles were also presented in two frozen noises ranging from 10 to 24 000 Hz. To generate these noises, recordings were performed in the colony room where a large group of guinea pigs were housed (30-40; 2-4 animals/cage). Several 4-seconds of audio recordings were added up to generate the “chorus noise”, which power spectrum was computed using the Fourier transform. This spectrum was then used to shape the spectrum of a white Gaussian noise. The resulting vocalization-shaped stationary noise therefore matched the “chorus-noise” audio spectrum. Figure 1A displays the overall envelopes of the four whistles in the vocalization-shaped stationary noise (third panel) and in the chorus noise (fourth panel) with a signal-to-noise ratio (SNR) of +10, 0 and −10 dB.

### Surgical procedures

All animals were anesthetized by an initial injection of urethane (1.2 g/kg, i.p.) supplemented by additional doses of urethane (0.5 g/kg, i.p.) when reflex movements were observed after pinching the hind paw (usually 2-4 times during the recording session). A single dose of atropine sulfate (0.06mg/kg, s.c.) was given to reduce bronchial secretions and a small dose of buprenorphine was administrated (0.05mg/kg, s.c.) as urethane has no antalgic properties. After placing the animal in a stereotaxic frame, a craniotomy was performed and a local anesthetic (Xylocain 2%) was liberally injected in the wound.

For auditory cortex recordings (area A1 and VRB), a craniotomy was performed above the left temporal cortex. The dura above the auditory cortex was removed under binocular control and the cerebrospinal fluid was drained through the cisterna to prevent the occurrence of oedema. For the recordings in MGB, a craniotomy was performed on the most posterior part of the MGB (8mm posterior to Bregma) to reach the left auditory thalamus at a location where the MGB is mainly composed of its ventral, tonotopic, part (Redies et al., 1989, Edeline et al., 1999; Anderson et al., 2007; Wallace et al., 2007). For IC recordings, a craniotomy was performed above the IC and portions of the cortex were aspirated to expose the surface of the left IC (Malmierca et al., 1995, 1996; Rees et al., 1997). For CN recordings, after opening the skull above the right cerebellum, portions of the cerebellum were aspirated to expose the surface of the right CN (Paraouty et al., 2018).

After all surgeries, a pedestal in dental acrylic cement was built to allow an atraumatic fixation of the animal’s head during the recording session. The stereotaxic frame supporting the animal was placed in a sound-attenuating chamber (IAC, model AC1). At the end of the recording session, a lethal dose of Exagon (pentobarbital >200 mg/kg, i.p.) was administered to the animal.

### Recording procedures

Data from multi-unit recordings were collected in 5 auditory structures, the non-primary cortical area VRB, the primary cortical area A1, the medial geniculate body (MGB), the inferior colliculus (IC) and the cochlear nucleus (CN). In a given animal, neuronal recordings were only collected in one auditory structure.

Cortical extracellular recordings were obtained from arrays of 16 tungsten electrodes (TDT, TuckerDavis Technologies; ø: 33 μm, <1 MΩ) composed of two rows of 8 electrodes separated by 1000 μm (350 μm between electrodes of the same row). A silver wire, used as ground, was inserted between the temporal bone and the dura mater on the contralateral side. The location of the primary auditory cortex was estimated based on the pattern of vasculature observed in previous studies (Wallace et al., 2000; Gaucher et al., 2013, 2020; Gaucher and Edeline, 2015). The non-primary cortical area VRB was located ventral to A1 and distinguished because of its long latencies to pure tones (Rutkowski et al., 2002; Grimsley et al., 2012). For each experiment, the position of the electrode array was set in such a way that the two rows of eight electrodes sample neurons responding from low to high frequency when progressing in the rostro-caudal direction [see examples in Figure 1 of Gaucher et al., (2012) and in Figure 6A of Occelli et al., (2016)].

In the MGB, IC and CN, the recordings were obtained using 16 channel multi-electrode arrays (NeuroNexus) composed of one shank (10 mm) of 16 electrodes spaced by 110 μm and with conductive site areas of 177μm^2^. The electrodes were advanced vertically (for MGB and IC) or with a 40° angle (for CN) until evoked responses to pure tones could be detected on at least 10 electrodes.

All thalamic recordings were from the ventral part of MGB (see above surgical procedures) and all displayed latencies < 9ms. At the collicular level, we distinguished the lemniscal and non-lemniscal divisions of IC based on depth and the latencies of pure tone responses. We excluded the most superficial recordings (until a depth of 1500μm) and those exhibiting latency >= 20ms in an attempt to select recordings from the central nucleus of IC (CNIC). At the level of the cochlear nucleus, the recordings were collected from both the dorsal and ventral divisions.

The raw signal was amplified 10,000 times (TDT Medusa). It was then processed by an RX5 multichannel data acquisition system (TDT). The signal collected from each electrode was filtered (610-10000 Hz) to extract multi-unit activity (MUA). The trigger level was set for each electrode to select the largest action potentials from the signal. On-line and off-line examination of the waveforms suggests that the MUA collected here was made of action potentials generated by a few neurons at the vicinity of the electrode. However, as we did not used tetrodes, the result of several clustering algorithms (Pouzat et al., 2002; Quiroga et al., 2004; Franke et al., 2015) based on spike waveform analyses were not reliable enough to isolate single units with good confidence. Although these are not direct proofs, the fact that the electrodes were of similar impedance (0.5-1 MOhm) and that the spike amplitudes had similar values (100-300 μV) for the cortical and the subcortical recordings, were two indications suggesting that the cluster recordings obtained in each structure included a similar number of neurons. Even if a similar number of neurons were recorded in the different structures, we cannot discard the possibility that the homogeneity of the multi-unit recordings differ between structures. By collecting several hundreds of recordings in each structure, these potential differences in homogeneity should be attenuated in the present study.

### Simulations of auditory nerve fiber responses

A computational model of auditory nerve fiber responses was used to assess whether the envelope-tracking properties measured in the central auditory system could be a mere consequence of the processing taking place at peripheral levels. For this purpose, an improved version of a well-established and widely-used model of the auditory periphery was used (Bruce, Erfani & Zilany, 2018; earlier versions of the model include Zilany & Bruce 2007 and Zilany, Bruce & Carney 2014). This model provides a phenomenological description of the major functional stages of the auditory periphery, from the middle ear up to the auditory nerve (Osses et al., in press). The implementation used in the present study is available as the routine ‘bruce2018 ‘within the AMT toolbox (v1.0) for MATLAB (Majdak et al., in press).

In order to make the simulated data as comparable as possible to the neuronal responses collected in the electrophysiological experiments, the distribution of cochlear center frequencies was chosen to be similar to the best frequencies obtained from the CN data. Default parameters were used for the later stages of the model. For each cochlear channel, five auditory-nerve fibers were simulated with different spontaneous rates (SR): 1 low-SR fiber (SR = 0.1 spikes/s), 1 medium-SR fiber (SR = 4 spikes/s) and 3 high-SR fibers (SR = 100 spikes/s). The outcome of the model corresponds to the aggregated responses of these 5 simulated auditory nerve fibers (sANF) in an attempt (i) to keep the physiological ratio between low, medium and high threshold fibers and (ii) to be close from the MUA collected in the auditory structures.

The responses to twenty repetitions of each vocalization in the clean and degraded conditions were simulated and analyzed in the same way as recorded data.

### Experimental protocol

As inserting an array of 16 electrodes in a brain structure almost systematically induces a deformation of this structure, a 30-minutes recovering time lapse was allowed for the structure to return to its initial shape, then the array was slowly lowered. Tests based on measures of time-frequency response profiles (TFRPs) were used to assess the quality of our recordings and to adjust electrodes’ depth. For auditory cortex recordings (A1 and VRB), the recording depth was 500-1000 μm, which corresponds to layer III and the upper part of layer IV according to Wallace and Palmer (2008). For thalamic recordings, the NeuroNexus probe was lowered about 7mm below pia before the first responses to pure tones were detected. For the collicular and cochlear nucleus recordings, the NeuroNexus probe was visually inserted in the structure and after a 15 minutes stabilization period, auditory stimuli were presented.

When a clear frequency tuning was obtained for at least 10 of the 16 electrodes, the stability of the tuning was assessed: we required that the recorded neurons displayed at least three successive similar TFRPs (each lasting 6 minutes) before starting the protocol. When the stability was satisfactory, the protocol was started by presenting the acoustic stimuli in the following order: We first presented the four whistles at 75 dB SPL in their original versions (in quiet), then the chorus and the vocalization-shaped stationary noises were presented at 75 dB SPL followed by the masked vocalizations presented against the chorus then against the vocalization-shaped stationary noise at 65, 75 and 85 dB SPL. Thus, the level of the original vocalizations was kept constant (75 dB SPL), and the noise level was increased (65, 75 and 85 dB SPL). In all cases, each vocalization was repeated 20 times. Presentation of this entire stimulus set lasted 45 minutes. The protocol was restarted either after moving the electrode arrays on the cortical map or after lowering the electrode by at least 300 μm for subcortical structures.

### Behavioral Go/No-Go discrimination task

Nine eight-weeks old C57Bl/6J mice were water-deprived and trained daily for 200–300 trials in a Go/No-Go task involving two of the guinea pig whistles (W1 and W3), one signaling the reward (the S+) and the other not (the S-). The training procedures were similar to those described in previous studies (Deneux et al., 2016; Ceballo et al., 2019). Water-deprived mice (33 ml/g per day) were head-fixed and held in a plastic tube on aluminium foil. Mice first performed 1-3 habituation sessions to learn to obtain a water reward (~5 μl) by licking on a spout over a threshold after the positive stimulus S+. Licks were detected by changes in resistance between the aluminium foil and the water spout. After habituation, the fraction of collected rewards was ~80%.

The learning protocol then started in which mice also received a non-rewarded, the S- for which they had to decrease licking below threshold to avoid an 8s time-out. One of the two whistles (the S+ or the S-) was presented every 10-20 s (uniform distribution) followed by a 1s test period during which the mouse had to produce at least one lick on a stainless steel water spout to receive the 5 μl water drop. Positive and negative stimuli were played in a pseudorandom order with the constraint that exactly 4 positive and 4 negative sounds must be played every 8 trials.

Once a mouse show at least 80% of correct discrimination between the S+ and the S- for two successive days in the original condition, it was trained in noisy conditions first with the stationary noise (successively at +10, 0 and −10 dB SNR) then with the chorus noise (successively at +10, 0 and −10 dB SNR). Each mouse had to perform at least one day at 80% in a given SNR to be tested on the following day at a lower SNR. Behavioral analyses were all automated thus no animal randomization or experimenter blinding was used.

## Data analysis

All the analyses were performed on MATLAB 2021 (MathWorks).

### Quantification of responses to pure tones

The TFRP were obtained by constructing post-stimulus time histograms for each frequency with 1 ms time bins. The firing rate evoked by each frequency was quantified by summing all the action potentials from the tone onset up to 100 ms after this onset. Thus, TFRP were matrices of 100 bins in abscissa (time) multiplied by 129 bins in ordinate (frequency). All TFRPs were smoothed with a uniform 5×5 bin window. For each TFRP, the Best Frequency (BF) was defined as the frequency at which the highest firing rate was recorded. Peaks of significant response were automatically identified using the following procedure: A positive peak in the TFRP was defined as a contour of firing rate above the average level of the baseline activity plus six times the standard deviation of the baseline activity. Recordings without significant peak of responses or with inhibitory responses were excluded from the data analyses.

### Quantification of the envelope tracking

We first filtered all the overall envelopes (original, vocoded and noisy vocalizations) using a bank of 35 gammatone filters with center frequencies uniformly spaced along a guinea pig - adapted ERB (equivalent rectangular bandwidth) scale ranging from 20 to 30 000 Hz. Four ranges of amplitude modulation (AM) were investigated: the very low (VL, < 4 Hz), low (L, < 20 Hz), middle (M, between 20 and 100 Hz) and high (H, between 100 and 200 Hz) AM ranges. For all the AM filtering, we used Butterworth filters at −6 dB per octave. Second, the envelopes were downsampled to a resolution of 1 ms to match the sampling rate of the PSTHs. Finally, we applied a half-wave rectification followed by a normalization with the corresponding RMS value.

The neuronal responses (i.e. the PSTHs) were also filtered with the same four frequency bands as the envelopes followed by a normalization with the corresponding RMS value. The rationale for this filtering step was that we wanted to isolate and quantify the correspondence between particular temporal varying aspects of the stimuli and PSTHs.

Next, we performed normalized cross-correlations between the filtered envelopes and PSTHs for each AM range. We selected seven gammatones, as a trade-off between accurately representing the envelopes along the audio spectrum and minimizing redundancy between envelopes. Maximal values in the correlograms were automatically detected in each structure to account for propagation delays in the auditory system. The lags were selected according to the distributions of the latencies obtained in response to pure tones at 75 dB SPL. The different lags identified were: 1-10ms for CN, 5-20ms for CNIC, 6-15ms for MGv, 9-30ms for A1 and 9-40ms for VRB.

#### Evaluation of the correlation significance by shuffling the evoked activity

It is known that significant correlation between neuronal events and sensory stimuli can be obtained by chance (see for review Harris 2020). Therefore, it was crucial to run drastic controls to ensure that the correlations detected here did not result from spurious correlations.

To determine a significance threshold for the correlation, we shuffled only the evoked activity in the original condition on a time-scale of 1 ms, in order to preserve the global shape of the whole response (i.e., the four response peaks due to the starting of each stimulus). The obtained shuffled PSTHs were then processed using the same procedure as for unshuffled PSTHs: filtering in the four AM ranges and half-wave rectification followed by a normalization with the corresponding RMS value. Then, we computed the cross-correlation (RRandom) between each shuffled PSTH and each envelope. We performed this procedure 1000 times and set, for each correlation value PSTH-E, a significant threshold of the R value that is the mean of the R_Random_ values plus two fold the standard deviations (μ R_Random_ ± 2σ). Based upon this criterion, percentages of recordings were discarded in each structure and for each AM range: in VRB, 42%, 30%, 10%, 47% of recordings were discarded in the VL, L, M and H range respectively; In A1, 52%, 51%, 38%, 73% of recordings were discarded in the VL, L, M and H range respectively; In MGv, 32%, 61%, 49%, 63% of recordings were discarded in the VL, L, M and H range respectively; in CNIC, 52%, 29%, 26%, 33% of recordings were discarded in the VL, L, M and H range respectively; in CN, 49%, 33%, 43%, 50% of recordings were discarded in the VL, L, M and H range respectively; in sANF, 21%, 35%, 86%, 77% of recordings were discarded in the VL, L, M and H range respectively.

### Quantification of mutual information from the responses to vocalizations

The method developed by Schnupp and colleagues (2006) was used to quantify the amount of information contained in the responses to vocalizations obtained with natural, vocoded or noisy stimuli. This method allows quantifying how well the vocalization’s identity can be inferred from neuronal responses. Neuronal responses were represented using different time scales ranging from the duration of the whole response (total spike count) to a 1-ms precision (precise temporal patterns), which allows analyzing how much the spike timing contributes to the information. As this method is exhaustively described in Schnupp and colleagues (2006) and in Gaucher and colleagues (2013a), we only present below the main principles.

The method relies on a pattern-recognition algorithm that is designed to “guess which stimulus evoked a particular response pattern” (Schnupp et al., 2006) by going through the following steps: From all the responses of a subcortical or cortical site to the different stimuli, a single response (test pattern) is extracted and represented as a PSTH with a given bin size. Then, a mean response pattern is computed from the remaining responses for each stimulus class. The test pattern is then assigned to the stimulus class of the closest mean response pattern. This operation is repeated for all the responses, generating a confusion matrix where each response is assigned to a given stimulus class. From this confusion matrix, the Mutual Information (MI) is given by Shannon’s formula:

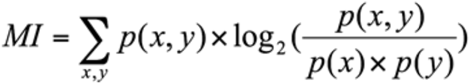

where x and y are the rows and columns of the confusion matrix, or in other words, the values taken by the random variables “presented stimulus class” and “assigned stimulus class”.

In our case, we used responses to the four whistles and selected the first 280 ms of these responses to work on spike trains of exactly the same duration (the shortest whistle being 280 ms long). In a scenario where the responses do not carry information, the assignments of each response to a mean response pattern is equivalent to chance level (here 0.25 because we used 4 different stimuli and each stimulus was presented the same number of times) and the MI would be close to zero. In the opposite case, when responses are very different between stimulus classes and very similar within a stimulus class, the confusion matrix would be diagonal and the mutual information would tend to log2(4) = 2 bits. This algorithm was applied with different bin sizes ranging from 1 to 280 ms (see figure 2B in Souffi and colleagues (2020) for the evolution of MI with temporal precisions ranging from 1 to 40 ms). The value of 8 ms was selected for the data analysis because in each structure the MI reached its maximum at this value of temporal precision.

The MI estimates are subject to non-negligible positive sampling biases. Therefore, as in Schnupp and colleagues (2006), we estimated the expected size of this bias by calculating MI values for “shuffled” data, in which the response patterns were randomly reassigned to stimulus classes. The shuffling was repeated 100 times, resulting in 100 MI estimates of the bias (MIbias). These MI_bias_ estimates are then used as estimators for the computation of the statistical significance of the MI estimate for the real (unshuffled) datasets: the real estimate is considered as significant if its value is statistically different from the distribution of MI_bias_ shuffled estimates. Significant MI estimates were computed for MI calculated from neuronal responses under one electrode. The range of MI_bias_ values was very similar between brain structure: depending on the conditions (original, vocoded and noisy vocalizations), it was from 0.102 to 0.107 bits in the CN, from 0.107 to 0.110 bits in the IC, from 0.105 to 0.114 bits in the MGB, 0.107 to 0.111 bits in the A1 and from 0.106 to 0.116 bits in VRB. There was no significant difference between the mean values of MI_bias_ in the different structures (unpaired t-test, all p>0.25).

### Quantification of acoustic envelope similarity

For each acoustic condition and each AM range, we estimated the acoustic similarity between each pair of stimuli as the correlation between their envelopes across the seven selected gammatones. Then, we averaged the six correlation values (related to all possible combinations with the four stimuli) to obtain an estimate of the similarity between the four stimuli for each condition (original, vocoding and noisy conditions) and each AM range (see Fig. 8A, dark lines). In order to confirm that there is no bias in our gammatone selection, we carried the same analysis on the output of the 35 gammatones and obtained similar results (see Fig. 8A, light lines).

### Statistical analysis

We used an analysis of variance (ANOVA) for multiple factors to reveal the main effects in the whole data set (vocoding conditions: three levels, masking noise conditions: three levels; auditory structures: six levels; AM ranges: four levels). Post-hoc pairwise tests were performed between the original condition and the vocoding or noisy conditions, or between structures to assess the significance of the multiple comparisons. They were corrected for multiple comparisons using Bonferroni corrections and were considered as significant if their p-value was below 0.05.

## Acknowledgments

JME was supported by grants from the French Agence Nationale de la Recherche (ANR) (ANR-14-CE30-0019-01). SS was supported by the Fondation pour la Recherche Médicale (FRM) grant number ECO20160736099 and by the Entendre Foundation. We thank Nihaad Paraouty for training us on the cochlear nucleus surgery. We thank Prf. Christian Lorenzi and Prf. Shihab Shamma for helpful comments on a previous version of the MS, and are particularly grateful to Dr. Virginie van Wassenhove for suggesting several improvements of the MS. We also wish to thank Céline Dubois, Mélanie Dumont and Aurélie Bonilla, for taking care of the guinea-pig colony.

## Competing Interests statement

The authors declare no competing financial interests.

## Notes

### Competing Interest Statement

The authors have declared no competing interest.

